# Heterologous expression in *Saccharomyces* and *Chlamydomonas* reveals host-dependent activity of *Brassica juncea* fatty acid elongase1 isozymes

**DOI:** 10.1101/2025.05.20.655142

**Authors:** Neelesh Patra, Susamoy Sarkar, Mrinal K. Maiti

## Abstract

The *fatty acid elongase1* (*FAE1*) genes of tetraploid *Brassica juncea* are the key determinant of high erucic acid (EA, C22:1) accumulation in its seed oil. While our previous work demonstrated near-zero EA content in mustard oil via CRISPR/Cas9 knockout of the two homeoalleles, *BjFAE1.1* and *BjFAE1.2*; the contributory function of each isozymes towards EA biosynthesis remains elusive. This study investigated the heterologous expression of *BjFAE1.1* and *BjFAE1.2* from high EA *B. juncea* cultivar JD6 in two metabolically distinct eukaryotic microbial hosts: the green microalga *Chlamydomonas reinhardtii* and the budding yeast *Saccharomyces cerevisiae*. Despite confirmed protein expression, neither BjFAE1 isozyme produced detectable C20:1 or C22:1 very-long-chain fatty acids (VLCFAs) in transgenic lines of *C. reinhardtii*. In contrast, expression in *S. cerevisiae* resulted in significant *de novo* biosynthesis of VLCFAs, C20:1 (∼9-11%) and C22:1 (∼17-19%), confirming their enzymatic activity as functional β-ketoacyl-CoA synthase. Substrate feeding experiments in yeast further validated their capability to elongate oleoyl-CoA (C18:1-CoA) to erucoyl-CoA (C22:1-CoA) via eicosenoyl-CoA (C20:1-CoA), with BjFAE1.1 showing slightly higher activity, as indicated by the enhanced VLCFAs accumulation. These findings highlight the critical influence of the heterologous host’s cellular environment on the enzyme functionality of plant genes involved in lipid metabolism, underscoring challenges for VLCFA production in microalgal platform.

## Introduction

Very-long-chain fatty acids (VLCFAs), typically defined as fatty acids (FAs) with acyl chain lengths of 20 carbons or more, are essential lipid molecules found across different domains of life. While eukaryotes utilize a conserved endoplasmic reticulum (ER)-localized elongase complex for their synthesis, prokaryotes employ diverse mechanisms; for instance, the Fatty Acid Synthase II (FAS-II) system generates unique VLCFAs like mycolic acids in *Mycobacterium* sp. (Marrakchi et al., 2014), whereas polyketide synthase-like enzymes are responsible for producing long-chain polyunsaturated fatty acids (PUFAs) in bacteria such as *Shewanella* sp. (Metz et al., 2001). In eukaryotes, VLCFAs serve critical functions as components of cellular membranes (e.g., sphingolipids, glycosylphosphatidylinositol anchors) and as precursors for vital structural lipids like cuticular waxes and suberin in plants, as well as storage lipids in seeds of certain plant species (Baud and Lepiniec, 2009; Li-Beisson et al., 2010). The biosynthesis of VLCFAs in plants occurs through sequential cycles of two-carbon chain elongation mediated by the ER membrane-bound multi-enzyme fatty acid elongase (FAE) complex (Baud and Lepiniec, 2009; Li-Beisson et al., 2010). The FAE complex catalyzes four core enzymatic reactions: condensation, reduction, dehydration, and a second reduction, for each two-carbon chain elongation (Baud and Lepiniec, 2009). The initial condensation step, joining malonyl-CoA with a long-chain acyl-CoA substrate (typically C18 or longer), is performed by β-ketoacyl-CoA synthase (KCS) enzyme, and is considered the rate-limiting, substrate-specific reaction determining overall flux through the VLCFA pathway (Millar and Kunst, 1997). Crucially, the substrate specificity of the particular KCS isoform dictates the chain length of the FAs produced.

Most of the *Brassicaceae* family plants, including the important oilseed crop *Brassica juncea* (Indian mustard), accumulate high levels (40-50%) of the VLCFA erucic acid (EA, C22:1ω-9) in their seed oil (Blacklock and Jaworski, 2006; Gupta et al., 2004; Patra et al., 2025). This high EA content is valuable as industrial feedstocks for applications including lubricants, surfactants, polymers, and slip agents like erucamide (Leonard, 1996), but limits the oil’s food use due to health concerns and regulatory restrictions (Abdellatif and Vles, 1970). Therefore, developing alternative, sustainable platforms for EA biosynthesis, such as microbial bioproduction systems, are highly desirable to meet industrial demand without compromising the development of low-EA *B. juncea* cultivars for improved edible oil quality. The *fatty acid elongase 1* (*FAE1*) genes encode KCS enzymes that are responsible for initiating VLCFA synthesis through preferential elongation of oleoyl-CoA (C18:1-CoA) substrate to eicosenoyl-CoA (C20:1-CoA) and subsequently to erucoyl-CoA (C22:1-CoA) (Millar and Kunst, 1997; Blacklock and Jaworski, 2006). In allotetraploid *B. juncea*, two functional *FAE1* homeoalleles (*BjFAE1.1* and *BjFAE1.2*) contribute to the high-erucic acid (HEA) phenotype (Gill et al., 2021; Katavic et al., 1995). Our prior research utilizing CRISPR/Cas9 knockout (KO) technology provided definitive *in planta* evidence that the simultaneous loss of both *BjFAE1.1* and *BjFAE1.2* function is necessary to eliminate EA accumulation in HEA *B. juncea* seeds (Patra et al., 2025). This targeted disruption of both homeoalleles resulted in a near-complete elimination of EA (reduced to <0.5%) in the mature seed oil, concurrently increasing the proportion of nutritionally desirable other FAs, while preserving essential agronomic traits such as plant growth and overall oil content (Patra et al., 2025). This CRISPR/Cas9 reverse genetic approach established the central function of FAE1 enzymes in EA biosynthesis within the complex metabolic network of the developing seed tissues of *B. juncea*; however, it precluded biochemical characterization of the individual enzymes encoded by each homeoallele. A comprehensive biochemical understanding necessitates functional analysis of the isozymes in simplified, defined heterologous expression systems outside the native plant cell. Key knowledge gaps remain regarding their intrinsic catalytic capabilities when isolated from the plant cellular context and potential subtle functional differences between the two homeologs. Moreover, elucidating the influence of distinct host cellular environments on the functionality of plant lipid metabolic enzymes, such as KCS, is crucial for accurately predicting feasibility and optimizing the outcomes in heterologous microbial expression systems designed for metabolic engineering, an area where predictive understanding remains incomplete.

To address these gaps, we employed a comparative heterologous expression strategy using two well-characterized, yet metabolically distinct, single-celled eukaryotic model systems: the green microalga *Chlamydomonas reinhardtii* and the budding yeast *Saccharomyces cerevisiae*. Model eukaryotic microbial systems represent important sustainable platforms for metabolic engineering due to their genetic tractability, rapid growth, scale-up cultivation and established capacity for producing diverse oleochemicals and other valuable compounds (Adrio and Demain, 2014; Nielsen et al., 2013; Rosenberg et al., 2008). Their potential as cell factories allows for the abundant biosynthesis of specific target molecules in isolated controlled environment, bypassing agricultural limitations. Understanding the feasibility of producing VLCFAs like EA in these systems is crucial for developing alternative production strategies for industrial usage. *C. reinhardtii*, being a photosynthetic eukaryote, can be a sustainable microbial platform for EA production. Its lipid metabolism involves complex interplay between plastidial *de novo* FA biosynthesis and ER-based FA/lipid modification and trafficking pathways (Li-Beisson et al., 2019; Merchant et al., 2007), potentially offering a comparable substrate landscape and cellular environment similar to plant. Evaluating BjFAE1 function in *Chlamydomonas* allows assessment of enzyme adaptability in a single-cellular photosynthetic eukaryotic context relevant to algal biotechnology applications aimed at producing tailored lipids (Cahoon et al., 2007). The other model used in this study, *S. cerevisiae*, is frequently used for functional characterization of plant lipid metabolic enzymes (Lassner et al., 1996; Millar and Kunst, 1997; Schneiter et al., 2000) and possesses compatible ER-localized FAE complex components i.e., 3-ketoacyl-CoA reductase (KCR), 3-hydroxyacyl-CoA dehydratase (HCD), and trans-2,3-enoyl-CoA reductase (ECR). Its native elongation system, mediated by the Elo proteins (Elo1p, Elo2p, and Elo3p), primarily synthesizes acyl chains up to C18, although Elo3p specifically produces the C26 required for sphingolipid biosynthesis (Oh et al.,1997). Consequently, *S. cerevisiae* lacks endogenous production of C20:1 or C22:1 VLCFAs, making it highly suitable for validating the function of introduced plant KCS enzymes (Li-Beisson et al., 2010).

In the present study, the *B. juncea FAE1* homeoalleles, *BjFAE1.1* and *BjFAE1.2*, from the HEA cultivar JD6, were individually expressed in *C. reinhardtii* CC125 and *S. cerevisiae* INVSc1. Functional assessment of the expressed isozymes was performed by analyzing the total FA profiles of the recombinant strains for the production of the expected VLCFA products, C20:1 and C22:1, using gas chromatography-mass spectrometry (GC-MS). EA-precursor feeding experiments in yeast were conducted to confirm substrate specificity of BjFAE1 isozymes. This comparative analysis revealed distinct outcomes, with the consistent absence of VLCFAs in the microalgal host but clear functional enzyme activity yielding 25-30% combined C20:1 and C22:1 VLCFAs in the yeast system.

## Materials and methods

### Microbial strains and culture conditions

*Chlamydomonas reinhardtii* strain CC-125 (Chlamydomonas Resource Center; https://www.chlamycollection.org) were grown in tris-acetate-phosphate (TAP) medium at 25 °C with shaking at 120 rpm, under a 14-hr light (25 μM/m2/s) and 10-hr dark cycle. *Saccharomyces cerevisiae* strain INVSc1 (Invitrogen) was used for yeast expression studies. Yeast strains were routinely cultured in an orbital shaker at 30 °C and 120 rpm on yeast extract peptone dextrose (YEPD) medium (Himedia). For selection of transformants and gene expression studies, INVSc1 cells were grown in synthetic minimal dropout medium lacking uracil (SC-Ura) with either glucose and galactose, as reported earlier (Chattopadhyay et al., 2020)

### Construction of expression plasmids

The coding DNA sequences (1521 bp) of *B. juncea Fatty Acid Elongase1* homeoalleles, *BjFAE1.1* (GenBank accession no. <u>**PQ858696**</u>) and *BjFAE1.2* (GenBank accession no. <u>**PQ858697**</u>), were derived from the high-erucic acid *B. juncea* cultivar JD6, as previously characterized (Patra et al., 2025).

For expression in *C. reinhardtii*, the amplified *BjFAE1.1* and *BjFAE1.2* coding sequences without stop codon were independently cloned into the *C. reinhardtii* expression plasmid pOpt2_mVenus_Hyg (Wichmann et al., 2017), purchased from the Chlamydomonas Resource Center. For each *BjFAE1* gene, two versions of the construct were prepared, one with a 3’-end mVenus reporter gene tag [the gene of interest (GOI) without stop codon cloned using NdeI and BglII restriction enzymes (RE)], and the other without the tag (GOI cloned using NdeI and EcoRI RE).

For expression in *S. cerevisiae*, the *BjFAE1.1* and *BjFAE1.2* CDSs were cloned into the pYES2/CT plasmid (Invitrogen) using KpnI and EcoRI RE sites, placing them under the control of the galactose-inducible GAL1 promoter.

All recombinant plasmids were verified by appropriate RE digestion (**Figure S1, S3**) and Sanger sequencing to confirm the integrity and orientation of the inserts. All the primers used in this study are listed in **Table S1**.

### Transformation of *C. reinhardtii* and *S. cerevisiae*

*C. reinhardtii* strain CC-125 cells were transformed separately with the four prepared constructs (*BjFAE1.1*-mVenus, *BjFAE1.2*-mVenus, *BjFAE1.1, BjFAE1.2*) or an empty vector control (pOpt2_mVenus_Hyg without GOI) following the electroporation method described by Wittkopp (2018). Post-electroporation, cells were plated on TAP-agar selection medium containing 50 mg/L hygromycin (Duchefa Biochemie).

1. *cerevisiae* INVSc1 cells were transformed separately with pYES2/CT-*BjFAE1.1*, pYES2/CT-*BjFAE1.2*, or empty pYES2/CT following Invitrogen user manual.

### Verification of gene expression in transgenic *C. reinhardtii* lines

Transgenic CC-125 lines transformed with *BjFAE1.1*-mVenus and *BjFAE1.2*-mVenus fusion constructs were analyzed for mVenus fluorescent protein expression. Live cells were observed under an Olympus IX51 inverted fluorescence microscope equipped with a BUC4-140C colour cool CCD 1.4 MP digital camera. Image capture and processing were performed using TCapture PC software with appropriate filter sets for mVenus detection. For transformants generated with constructs lacking the mVenus tag, total RNA was extracted using the QIAGEN RNeasy mini kit. Quality and quantity were checked on a 1.5% agarose gel electrophoresis and Thermo Scientific NanoDrop 2000 equipment, respectively. Genomic DNA contamination was removed using DNase I (Sigma) treatment according to the manufacturer’s instructions. One-step reverse transcription PCR (RT-PCR) was performed using the QIAGEN OneStep RT-PCR Kit. *BjFAE1* expression was confirmed by the presence of a ∼165 bp amplicon, and the *Chlamydomonas* G-protein beta subunit-like polypeptide (CBLP) gene (∼460 bp amplicon) was used as an internal reference (Schloss, 1990). All the primers used in this study are listed in **Table S1**.

### Lipid extraction and fatty acid methyl ester (FAME) analysis by GC-MS

Total lipids were extracted from harvested *C. reinhardtii* cells (∼50 ml early stationary phase culture) and galactose-induced *S. cerevisiae* cells (∼50 mL, 48 h after of galactose induction cultures) using a Bligh and Dyer (1959) based method. Individual extracted samples were converted to FAMEs analyzed by using CLARUS 690 GC/MS instrument (Perkin Elmer) with Elite-5MS column following Patra et al., 2025.

### Substrate feeding experiment in *S. cerevisiae*

Transformed *S. cerevisiae* INVSc1 cells were grown in SC-Ura medium supplemented with 2% (w/v) glucose until mid-log phase. Gene expression was then induced by transferring cells to SC-Ura medium supplemented with 2% (w/v) or 4% (w/v) galactose. After 24 hours of induction, cultures were supplemented with oleic acid (C18:1ω9) and eicosenoic acid (C20:1ω9) separately to a final concentration of 50 µM. Stock solutions of FAs were prepared in DMSO and with a carrier like 0.1% BSA to aid solubilization. Control cultures received an equivalent volume of the carrier without any FA. The supplemented cultures were incubated for an additional 48 hours at 30 °C with shaking. Cells were then harvested, and total lipids were extracted and analyzed for FAME composition as described in section 2.5.

## Results

### Expression of BjFAE1.1 or BjFAE1.2 in *C. reinhardtii* does not alter FA profile

Transgenic *C. reinhardtii* lines successfully expressing either *BjFAE1.1* or *BjFAE1.2* from *B. juncea* cv. JD6 were generated and analyzed. Genetic constructs included versions of both with and without a C-terminal mVenus tag (**Figure 1a**) to facilitate expression verification and control for potential tag interference. Fluorescence microscopy readily confirmed the presence of the mVenus-tagged fusion proteins within the algal cells (**Figure S1**). For transformants generated using constructs lacking the mVenus tag, the expression of BjFAE1 was confirmed by reverse transcription PCR (RT-PCR) (**Figure S1**) using *Chlamydomonas* G-protein beta subunit-like polypeptide (CBLP) expression as internal reference (Schloss, 1990). Importantly, the introduction and expression of these plant elongase enzymes did not cause any apparent physiological burden; the generated transgenic algal lines displayed normal growth rates and cell morphology comparable to the untransformed wild type (WT) control strain CC125 under standard culture conditions, indicating no obvious adverse effects on algal cell viability.

**Figure 1.**
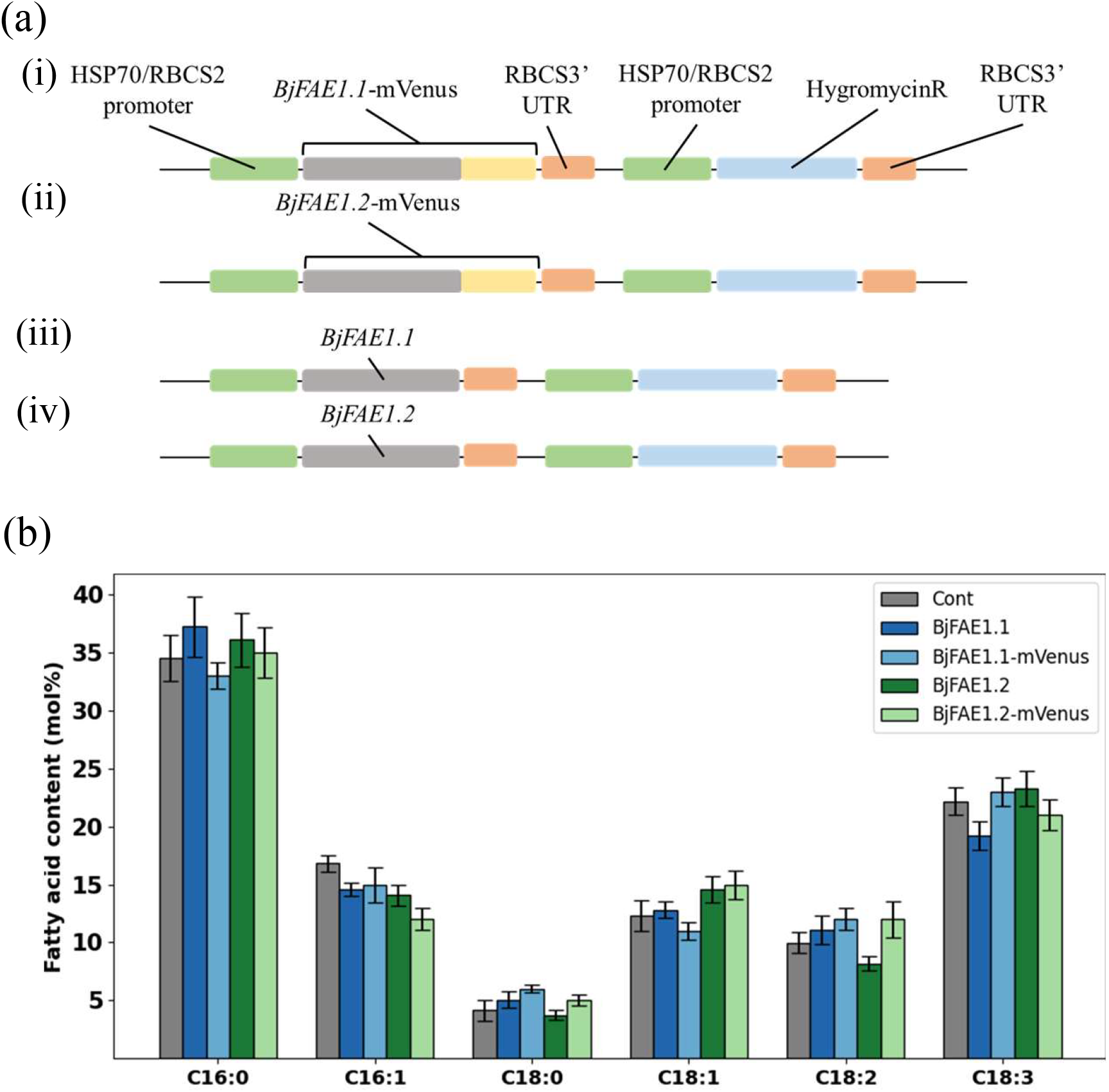
Heterologous expression of *BjFAE1* homeoalleles in *C. reinhardtii* strain CC-125 and FA profiling of stable transgenic algal lines. (**a**) Schematic representation of the expression constructs. (i) *BjFAE1.1*-mVenus, (ii) *BjFAE1.2*-mVenus, (iii) *BjFAE1.1* (untagged), and (iv) *BjFAE1.2* (untagged). (**b**) FA compositions (mol%) of individual transgenic algal lines along with empty vector transformed control, n=3 in each case. Values represent mean ± SE (n=3). Significant changes are calculated one-way ANOVA (analysis of variance) with post-hoc Tukey HSD (honestly significant difference) method. Since there is no significant change in FA composition between the control and the transgenic lines, the significance levels ‘*’ (p < 0.05) and ‘**’ (p < 0.01) are not shown in the bar graph.

Despite confirmed elongase protein expression, detailed GC-MS analysis of the total FA profiles showed that the transgenic *C. reinhardtii* lines (expressing either tagged or untagged BjFAE1) exhibit FA compositions essentially indistinguishable from the mock-transformed control strain (**Figure 1b**). The relative percentages of the major C16 and C18 FAs, including C16:0 (∼34-37%), C16:1 (∼14-17%), C18:0 (∼4-5%), C18:1 (∼12-15%), C18:2 (∼8-11%), and C18:3 (∼19-23%), showed no statistically significant differences between the control and the BjFAE1.1 or BjFAE1.2 expressing lines (**Figure 1b**). Careful examination of the chromatograms revealed no novel peaks or significant quantitative shifts in existing C16 and C18 FAs, which were typical of WT *C. reinhardtii*. Specifically, for BjFAE1 isoenzyme function, no new peaks corresponding to the expected VLCFA products, such as eicosenoic acid (C20:1) and/or EA, were detected above background levels in any of the analyzed transgenic algal lines (**Figure S2**). This lack of detectable product formation indicated that neither BjFAE1.1 nor BjFAE1.2 exhibited measurable elongase activity within the *C. reinhardtii* cellular environment under the conditions tested.

### Expression of BjFAE1.1 or BjFAE1.2 in *S. cerevisiae* leads to production of C20:1 and C22:1 FAs

In contrast to the unexpected results observed in the microalga *C. reinhardtii* system, both BjFAE1 isozymes demonstrated clear enzymatic activity when individually expressed from chimeric constructs (**Figure 2a**) in *S. cerevisiae* (strain INVSc1) upon galactose induction. Control INVSc1 cells harboring empty pYES2/CT vector exhibited a typical yeast FA profile under the standard induction conditions, dominated by C16 and C18 saturated and monounsaturated FAs (palmitic, palmitoleic, stearic, and oleic acids) (**Figure 2b, S4**). As expected, and consistent with the known specificity of native yeast Elo proteins, no significant accumulation of FAs longer than C18 was observed in these control yeast cells. Feeding experiments confirmed efficient uptake of exogenous FAs; supplementation with 50 µM C18:1 increased cellular C18:1 level (∼34.8%), while feeding with 50 µM C20:1 resulted in its clear detection (∼12.7%), but neither feeding condition led to the production of C20:1 or C22:1 in the control yeast strain (**Figure 2c**).

**Figure 2.**
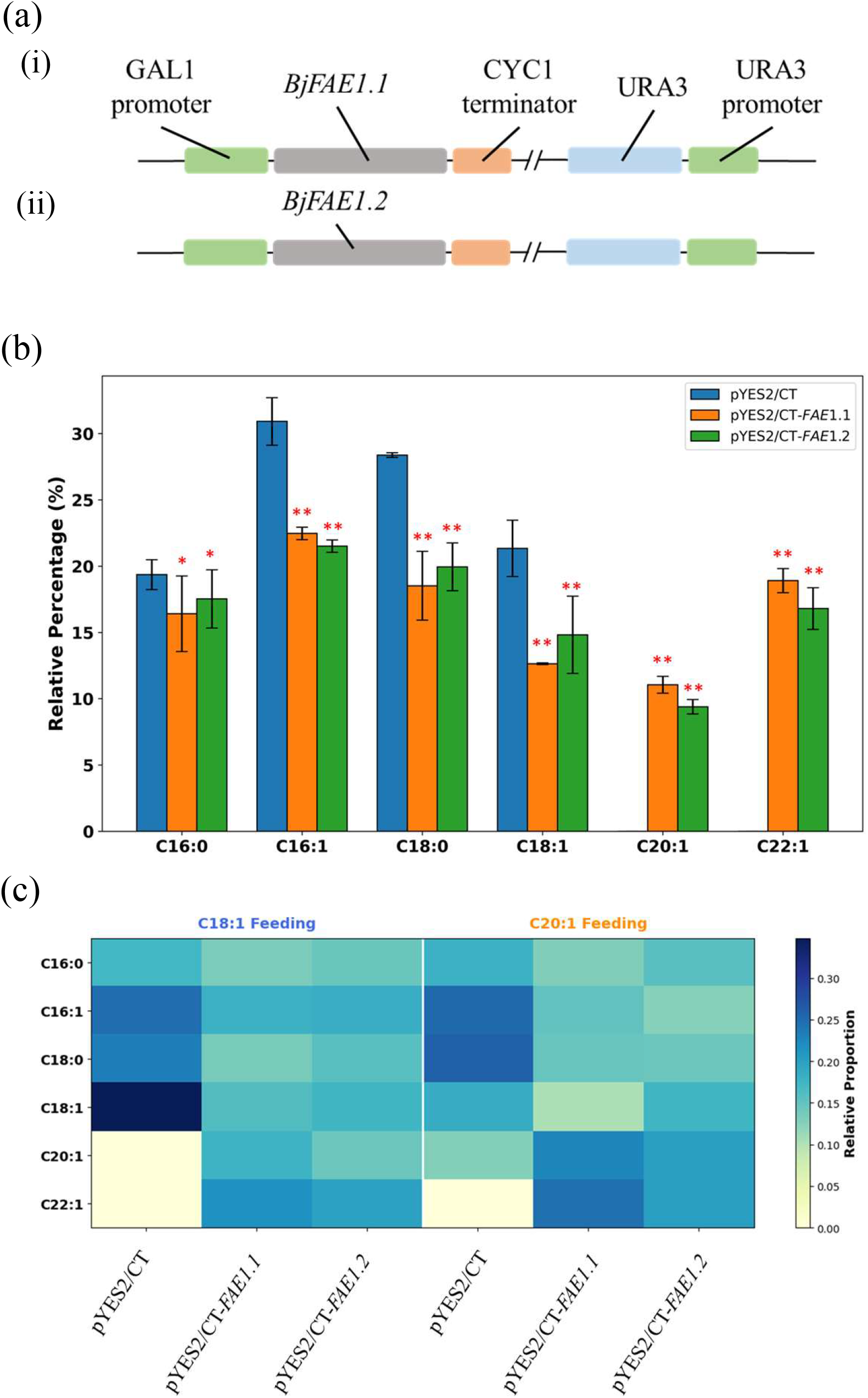
Heterologous expression of *BjFAE1* homeoalleles in *S. cerevisiae* INVSc1 strain and FA profiling of transgenic yeast lines under normal culture condition and FA-fed conditions. (**a**) Schematic representation of the expression constructs. (**i**) *BjFAE1.1* and (**ii**) *BjFAE1.2*, (**b**) FA compositions (mol%) of individual transgenic yeast lines along with empty vector transformed control, n=3 in each case. Values represent mean ± SE (n=3). Significant changes are calculated one-way ANOVA (analysis of variance) with post-hoc Tukey HSD (honestly significant difference) method. The significance levels are marked with ‘*’ (p < 0.05) and ‘**’ (p < 0.01) based on comparison with empty vector transformed control. (**c**) Heatmap showing the relative proportions of major FAs in transgenic yeast lines after feeding with either 50 µM C18:1 or 50 µM C20:1. Colour intensity represents the relative proportion of each FA, as indicated by the colour bar.

However, expression of *BjFAE1.1* chimeric construct (**Figure 2a**) under standard conditions resulted in the robust *de novo* production of both C20:1 (∼11.1%) and C22:1 (∼18.9%), unequivocally confirming functional enzyme activity (**Figure 2b, S4**). This synthesis of novel VLCFAs (totalling ∼30.0%) was accompanied by a significant reduction in the endogenous C18:1 precursor pool (to ∼12.6%), providing strong evidence for C18:1 utilization to produce VLCFAs. The pathway’s responsiveness to substrate availability was demonstrated when supplementation with 50 µM C18:1 increased total VLCFA levels (to ∼39.3%), particularly C20:1 (to ∼17.7%) and C22:1 (to ∼21.6%). Crucially, feeding with 50 µM C20:1 dramatically boosted total VLCFA production (to ∼47.1%), primarily by increasing both C20:1 (to ∼22.7%) and C22:1 (to ∼24.5%), confirming C20:1 as a key intermediate efficiently converted by BjFAE1.1 (**Figure 2c**).

Similarly, the expression of *BjFAE1.2* construct (**Figure 2a**) under the normal conditions yielded readily detectable levels of C20:1 (∼9.4%) and C22:1 (∼16.8%), totalling ∼26.2% VLCFAs, confirming its activity (**Figure 2b, S4**). Supplementation with C18:1 increased total VLCFA production (to ∼34.1%), elevating both C20:1 (to ∼14.4%) and C22:1 (to ∼19.7%). Feeding with C20:1 substantially elevated total VLCFA levels (to ∼40.0%), significantly increasing both C20:1 (to ∼20.0%) and C22:1 (to ∼20.0%), validating efficient catalysis of the second elongation step by BjFAE1.2 as well (**Figure 2c**).

These data indicate that both the homeoalleles of *BjFAE1* encode functionally active KCS isozymes capable of efficiently elongating C18:1 acyl substrates within the *S. cerevisiae* cellular context. Notably, BjFAE1.1 expression consistently yielded slightly higher levels of both C20:1 and C22:1 compared to BjFAE1.2, suggesting subtle differences in their specific activity or stability in this heterologous system. Furthermore, the significant production of these new VLCFAs in BjFAE1 transformed yeast was accompanied by an expected decrease in the relative percentage of C18:1, the precursor substrate of BjFAE1 compared to the empty vector control, strongly supporting the proposed biochemical function and substrate utilization. The substrate feeding experiments validated the C18:1 to C20:1 to C22:1 conversion pathway mediated by both the isozymes *in vivo*.

## Discussion

This study elucidates functional characteristics of the *B. juncea* FAE1 homeoalleles, *BjFAE1.1* and *BjFAE1.2*, through an examination of their activity in two evolutionarily distant heterologous eukaryotic hosts. Subsequent to our prior demonstration of their essential role *in planta* for EA biosynthesis (Patra et al., 2025), we explored the utility of these *BjFAE1* genes in the microalga *C. reinhardtii*, a well-established microbial platform for industrial bioproducts (Leon et al., 2013; Rosenberg et al., 2008). Surprisingly, these gene products (elongase isozymes) failed to yield detectable EA or any VLCFA when expressed in *C. reinhardtii*, despite confirmed protein expression. However, the same genes are expressed as functional enzymes capable of producing VLCFAs in the yeast *S. cerevisiae*, another well-known microbial platform. This divergence in outcomes highlights the significant influence of the host cellular environment on the activity of lipid metabolic enzymes and underscores potential constraints in transferring complex biosynthetic pathways between distinct eukaryotic organisms. The complete lack of detectable C20:1 or C22:1 accumulation in *C. reinhardtii* is unexpected, particularly given that the alga possesses the necessary genomic repertoire, including homologs for KCS, KCR, HCD, and ECR, suggesting a functional endogenous pathway for ER-based FA elongation (Merchant et al., 2007). The absence of enzyme product observed in our study, despite confirmation of heterologous protein expression, points towards significant bottlenecks within the wild-type algal host under the tested conditions. Several potential factors, individually or in combination, may account for this inactivity. Firstly, substrate limitation remains a strong possibility. Efficient transport of C18:1 from its primary synthesis site in the chloroplast to the ER, followed by activation to C18:1-CoA, may be rate-limiting. Furthermore, within the ER, the introduced BjFAE1 likely faces strong competition for the available C18:1-CoA pool from highly active native pathways, which include desaturases producing abundant C18 PUFAs and acyltransferases incorporating C18:1 into major membrane lipids (e.g., diacylglyceryl-N,N,N-trimethylhomoserine) or storage triacylglycerides (TAGs) (Bellou et al., 2014; Cahoon et al., 2007; Dolch et al., 2017). This metabolic channelling could severely restrict substrate availability for the heterologous KCS. Although specific *C. reinhardtii* starchless mutants under nitrogen starvation can accumulate certain distinct VLCFAs (e.g., C20:0, C22:0), albeit in small proportions (Li et al., 2010; Work et al., 2010), this occurs under conditions of major metabolic perturbation. Our findings demonstrate that simply overexpressing a potent plant KCS like BjFAE1 in the wild-type background under standard conditions is insufficient to overcome the inherent metabolic partitioning and drive detectable synthesis of its characteristic C20:1 and C22:1 products. Secondly, incompatibility with the host machinery or cellular environment represents another potential impediment. Plant FA elongation occurs via ER membrane-bound multi-enzyme complexes (Haslam and Kunst, 2013; Lassner et al., 1996), and optimal function may depend on specific protein-protein interactions or lipid environments within the ER membrane. The *B. juncea* KCS might not efficiently interact or form productive associations with the endogenous *C. reinhardtii* KCR, HCD, or ECR partners, hindering the completion of the elongation cycle (Bellou et al., 2014; Trenkamp et al., 2004). Thirdly, inefficient downstream processing or product instability could prevent accumulation. Unlike developing oilseeds, which possess a natural metabolic sink for incorporating VLCFAs into TAGs and storing them in lipid bodies, *C. reinhardtii* lacks this specialized capacity. The algal acyltransferases involved in TAG synthesis may exhibit poor affinity for C20:1-CoA or C22:1-CoA (Bellou et al., 2014). Consequently, any C20:1/C22:1 produced might fail to be incorporated into stable lipid pools, potentially leading to feedback inhibition, channelling into other lipid classes, or rapid degradation via peroxisomal β-oxidation (Kong et al., 2017) or other regulatory turnover mechanisms (Siaut et al., 2011). On the other hand, the significant production of both C20:1 (∼9-11%) and C22:1 (∼17-19%) FAs in yeast *S. cerevisiae* expressing either BjFAE1.1 or BjFAE1.2 provides direct biochemical evidence of their identity as functional KCS enzymes. The observed C22:1 accumulation level demonstrates effective heterologous function and is comparable to levels reported for other *Brassicaceae* FAE1 enzymes expressed in yeast, such as *B. napus* FAE1 alleles (∼10-15% C22:1) (Zafar et al., 2020) or *Crambe abyssinica* FAE1 (Lassner et al., 1996), and notably higher than *Eruca vesicaria* FAE1 (∼2.5% C22:1) (Li et al., 2012). Our results confirm that the BjFAE1 enzymes efficiently utilize endogenous yeast C18:1-CoA precursors (reducing C18:1 from ∼21.3% in control to ∼12.6-14.8% in transformants) and functionally couple with the downstream KCR, HCD and ECR components of the native yeast FA elongation system (Li-Beisson et al., 2010; Schneiter et al., 2000). The substrate feeding experiments further illuminate the pathway dynamics. Supplementation with C18:1 increased total VLCFA accumulation for both isozymes (to ∼39.3% for BjFAE1.1 and ∼34.1% for BjFAE1.2), suggesting that endogenous C18:1 supply can be a limiting factor for maximizing flux. More definitively, feeding with the intermediate precursor C20:1 led to a dramatic increase in total VLCFA accumulation (to ∼47.1% for BjFAE1.1 and ∼40.0% for BjFAE1.2), primarily boosting C22:1 level for BjFAE1.1 (to ∼24.5%) and significantly increasing both C20:1 (to ∼20.0%) and C22:1 (to ∼20.0%) for BjFAE1.2. This confirms C20:1 as a key intermediate and demonstrates that both isozymes efficiently catalyze the second elongation step (C20:1-CoA to C22:1-CoA). Interestingly, even when fed with C18:1, the primary substrate, a substantial portion is converted through C20:1 to C22:1, and feeding C20:1 further enhances C22:1 production, indicating a strong channeling preference towards the final C22:1 product irrespective of the initial substrate provided in this system. Comparing these findings with heterologous expression studies of other FAE1 orthologs provides valuable context. The *Arabidopsis thaliana* FAE1 (AtFAE1), when expressed in yeast, also produces both C20:1 and C22:1 from C18:1 precursor (Millar and Kunst, 1997), but is often considered less efficient in the second elongation step (C20:1 to C22:1) compared to the FAE1 enzymes from high-EA *Brassica* species (Paul et al., 2006; Lassner et al., 1996).

Similarly, expression of FAE1 genes from *Brassica napus* in yeast typically results in substantial C22:1 accumulation (Zafar et al., 2020), consistent with their role in high-EA rapeseed. Our results for BjFAE1.1 and BjFAE1.2, showing significant conversion to C22:1 (C22:1/C20:1 ratio ∼1.7-1.8 under unfed conditions) and efficient utilization of fed C20:1, align well with the functional profile expected for FAE1 enzymes from a high-EA *Brassica* species, confirming their competence in catalyzing both elongation steps required for EA synthesis. The observation that BjFAE1.1 expression consistently resulted in slightly higher VLCFA levels than BjFAE1.2 is also in agreement with our previous finding in the endogenous host *B. juncea* where homozygous FAE1.1 single mutant lines (e1e1E2E2) had lesser EA content than homozygous FAE1.2 single mutant lines (E1E1e2e2) (Patra et al., 2025). While modest, this difference may indicate subtle variations in intrinsic catalytic efficiency, protein stability, or interaction with other components of the elongation complex, potentially reflecting functional divergence between the *B. juncea FAE1* homeoalleles originating from the *B. rapa* and *B. nigra* progenitor genomes (Gill et al., 2021). This successful functional reconstitution confirms the utility of *S. cerevisiae* as a model system for dissecting the activity of plant KCS enzymes. The contrasting results obtained in *Chlamydomonas* alga and *Saccharomyces* yeast clearly illustrate that the metabolic context of the host cell is critical for the successful heterologous expression of lipid biosynthetic enzymes. Yeast proved permissive for BjFAE1 activity, likely benefiting from conserved ER functions and robust TAG synthesis. Microalgae like *C. reinhardtii*, despite possessing the basic enzymatic machinery (Merchant et al., 2007), present considerable challenges due to their unique metabolic architecture and regulation (Bellou et al., 2014; Siaut et al., 2011). While other algae like *Nanochloropsis* sp. and diatoms have been successfully engineered to produce specific VLCPUFAs (Hamilton et al., 2016), and *Botryococcus* naturally produces long-chain hydrocarbons derived from VLCFAs (Metzger and Largeau, 2005), achieving targeted production of VLCFAs like C22:1 in *Chlamydomonas* necessitates a more comprehensive, systems-level approach. This could involve engineering substrate precursor pools, co-expression of compatible partner enzymes (KCR, HCD, ECR) potentially from the source plant, optimizing codon usage, and modifying downstream metabolic sinks (e.g., enhancing TAG incorporation (Fan et al., 2011; Fan et al., 2012)) or blocking competing pathways like β-oxidation (Huerlimann and Roessler, 2018; Kong et al., 2017).

## Conclusion

In conclusion, this research genetically and biochemically confirms *B. juncea* FAE1.1 and FAE1.2 as functional KCS isozymes producing C20:1 and C22:1 VLCFAs in *S. cerevisiae*. However, the marked inactivity observed in *C. reinhardtii*, despite the host possessing the necessary genetic framework, underscores critical host-dependent barriers potentially related to substrate availability, machinery compatibility, or metabolic regulation. These findings contribute to the understanding of plant FAE1 function and highlight the complex interplay between heterologous enzymes and host cellular environments, providing crucial knowledge towards rational design of future metabolic engineering strategies, particularly for the production of VLCFA in microalga *C. reinhardtii*.

## Acknowledgements and funding sources

NP acknowledges financial support from UGC (India) and SERB (India). We thank Dr. Prabir Kumar Bhattacharyya (Associate Professor, Department of Genetics and Plant Breeding, Bidhan Chandra Krishi Vishwavidyalaya, India) for providing the *B. juncea* JD6 cultivar seeds used in the study. We also gratefully acknowledge the use of central facilities at IIT Kharagpur, India. This research was funded by SERB (India) grant CRG/2022/001705 awarded to MKM.

## Author contributions

NP: Conceptualization, Investigation, Methodology, Data curation, Formal analysis, Writing-original draft, Writing-review & editing. SS: Yeast methodology, Writing-review & editing. MKM: Project administration, Resources, Supervision, Funding acquisition, Writing– review & editing.

## Supporting information

[**Figures S1** to **S4** and **Table S1** are provided as single .docx file]

**Figure S1** Preparation of *BjFAE1* expression constructs and verification of transgenic *Chlamydomonas reinhardtii* lines expressing *BjFAE1* homeoalleles.

**Figure S2** Representative GC-MS chromatograms of FA profile of transgenic *C. reinhardtii* lines.

**Figure S3** Restriction enzyme digestion confirming *BjFAE1* expression constructs used for *S. cerevisiae* transformation.

**Figure S4** Representative GC-MS chromatograms of FA profile of transgenic *S. cerevisiae*.

**Table S1** List of primers used in this study.

