## Supplementary figures and table for "Heterologous expression in *Saccharomyces* and *Chlamydomonas* reveals host-dependent activity of *Brassica juncea* fatty acid elongase1 isozymes"

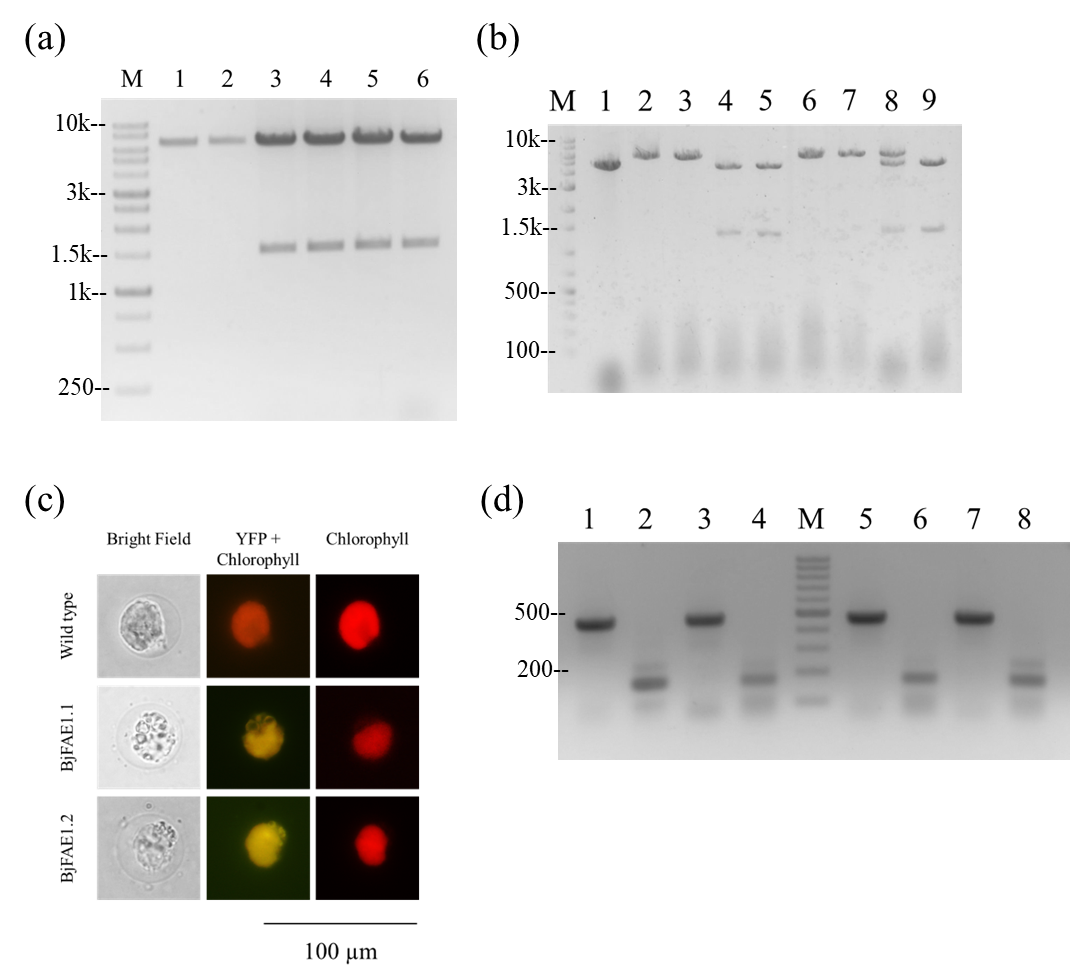

**Figure S1** Preparation of *BjFAE1* expression constructs and verification of transgenic *Chlamydomonas reinhardtii* lines expressing *BjFAE1* homeoalleles. (**a**) Restriction enzyme digestion confirming *BjFAE1*-mVenus tagged constructs used for algal transformation. Lane 1: control plasmid digested with NdeI; Lane 2: control plasmid digested with NdeI and BglII; Lanes 3 and 4: Plasmid from two *BjFAE1.1*-mVenus clones digested with NdeI and BglII, showing ~1.5 kb insert of *BjFAE1.1*; Lanes 5 and 6: Plasmid from two *BjFAE1.2*-mVenus clones digested with NdeI and BglII, showing ~1.5 kb insert of *BjFAE1.2*; M: DNA molecular weight marker in base pair (bp). **(b)** Agarose gel electrophoresis confirming untagged *BjFAE1* constructs. Lane 1: control undigested plasmid; Lanes 2 and 3: Undigested plasmids from two *BjFAE1.1* (untagged) clones; Lanes 4 and 5: Plasmids from the same *BjFAE1.1* clones digested with NdeI and EcoRI, showing the desired ~1.5 kb excised *BjFAE1.1* fragment; Lanes 6 and 7: Undigested plasmids from two *BjFAE1.2* (untagged) clones; Lanes 8 and 9: Plasmids from the same *BjFAE1.2* clones digested with NdeI and EcoRI, showing the desired ~1.5 kb excised *BjFAE1.2* fragment; M: DNA molecular weight marker in base pair (bp). **(c)** Fluorescence microscopy of transgenic algal cells expressing BjFAE1-mVenus fusion proteins. Representative images show bright-field, chlorophyll autofluorescence (red), and mVenus fluorescence (yellow). Wild-type cells are shown as a control. Scale bar = 100 µm. **(d)** RT-PCR products confirming *BjFAE1* transcript expression. Lanes 1, 3, 5, and 7 show the *CrCBLP* (constitutive housekeeping gene) amplicon of ~460 bp; Lanes 2 and 4: *BjFAE1.1* expression (amplicon size ~165 bp) from two independent transgenic algal lines developed with *BjFAE1.1* construct; Lanes 6 and 8: *BjFAE1.2* expression (amplicon size ~165 bp) from two independent transgenic algal lines developed with *BjFAE1.2* construct; M: 100-bp DNA ladder.

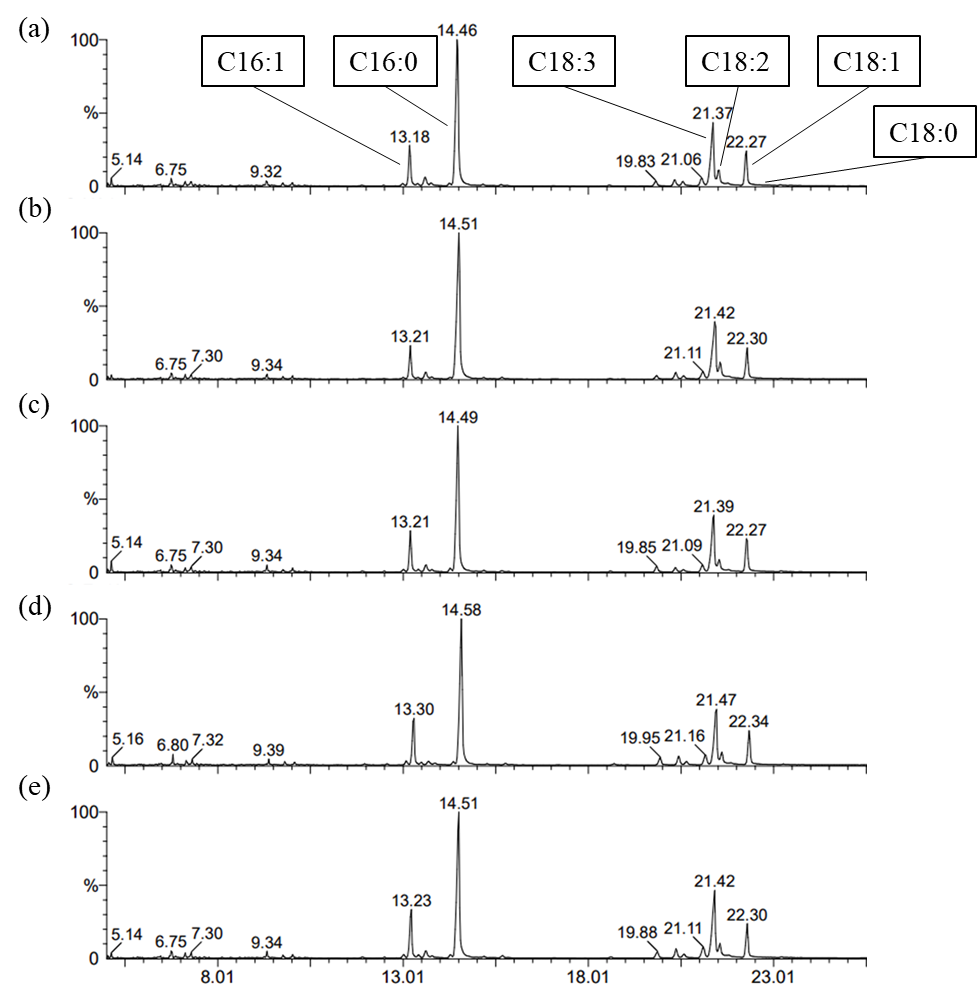

**Figure S2** Representative GC-MS chromatograms of FA profile of transgenic *C. reinhardtii* lines: (**a**) empty vector transformed control, (**b**) *BjFAE1.1*-mVenus, (**c**) *BjFAE1.2*-mVenus, (**d**) untagged *BjFAE1.1*, and (**e**) untagged *BjFAE1.2*.

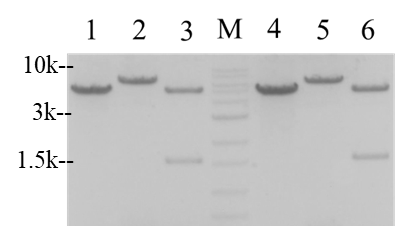

**Figure S3** Restriction enzyme digestion confirming *BjFAE1* expression constructs used for *S. cerevisiae* transformation. Lanes 1 and 4: Control undigested pYES2/CT plasmid; Lane 2: Undigested plasmid from one *BjFAE1.1* clone; Lane 3: Plasmid from the same *BjFAE1.1* clone digested with KpnI and EcoRI, showing the desired ~1.5 kb excised *BjFAE1.1* fragment; Lane 5: Undigested plasmid from one *BjFAE1.2* clone; Lane 6: Plasmid from the same *BjFAE1.2* clone digested with KpnI and EcoRI, showing the desired ~1.5 kb excised *BjFAE1.1* fragment; M: DNA molecular weight marker in base pair (bp).

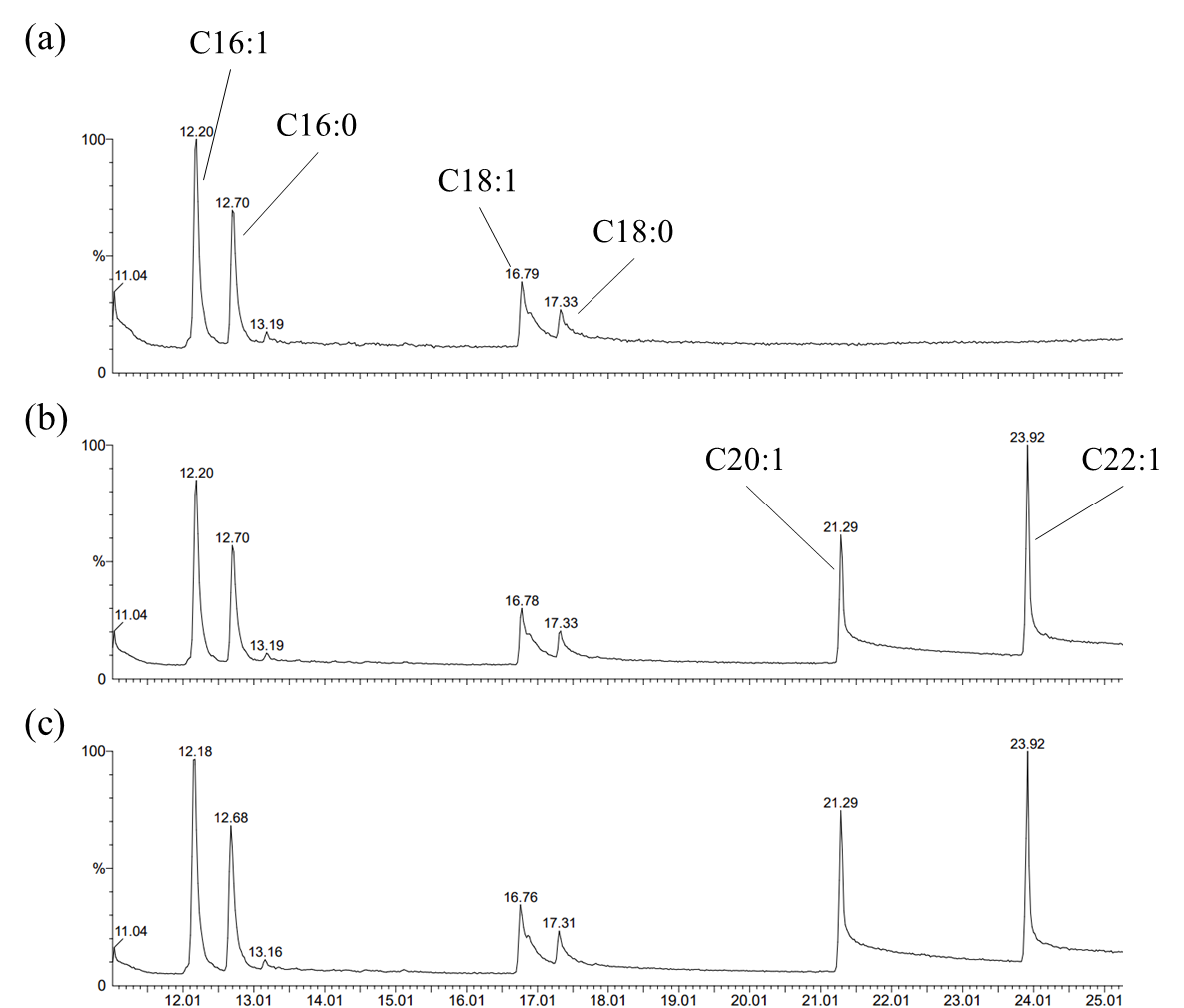

**Figure S4** Representative GC-MS chromatograms of FA profile of transgenic *S. cerevisiae*: (**a**) empty vector transformed control, (**b**) *BjFAE1.1* and (**c**) *BjFAE1.2*.

**Table S1** List of primers used in this study

| **Primer name** | **Sequence (5’-3’)** |
| --- | --- |
| BjFAE1_pOpt_NdeI_For | CCTGCATATGACGTCCATTAACGTAAAGCTCC |
| BjFAE1.1_pOpt_BglII_Rev | CCGAGATCTGGACCGACCGTTTTGGACAC |
| BjFAE1.1_pOpt_pYES_EcoRI_Rev | GGCGAATTCTTAGGACCGACCGTTTTGGACAC |
| BjFAE1_pYES_KpnI_For | GTCGGTACCAACACAATGTCTATGACGTCCATTAACGTAAAG |
| BjFAE1.1_pYES_EcoRI_Re | GGCGAATTCTTAGGACCGACCGTTTTGGACAC |
| BjFAE1.2_pOpt_BglII_Rev | CCGAGATCTGGACCGACCGTTTTGGACACGAAC |
| BjFAE1.2_pOpt_pYES_EcoRI_Rev | GGCGAATTCTTAGGACCGACCGTTTTGGACACGAAC |
| BjFAE1_ORT_For | GTGGCTTGACTTCTTGAGGA |
| BjFAE1.1_ORT_Rev | GTGTTTTTGAATAGATTCTC |
| BjFAE1.2_ORT_Rev | GTGTTCTCGAATAGATTTTT |
| CrCBLP_ORT_For | GACAACCGCCAGATCGTGTCGGGCTC |
| CrCBLP_ORT_Rev | CACGATGCTCTTGCTCTCCAGGTCCC |
